# Surveillance of migratory shorebirds and seabirds in 2024 in Australia reveals incursions of a diversity of low pathogenicity avian influenza viruses though not high pathogenicity avian influenza H5N1

**DOI:** 10.1101/2025.11.06.687052

**Authors:** Michelle Wille, Robyn Atkinson, Ian G. Barr, Alexander L. Bond, David Boyle, Maureen Christie, Yi-Mo Deng, Meagan L. Dewar, Tegan Douglas, Min He, Rosalind Jessop, Lindall R. Kidd, Hiske Klaassen, Jennifer Lavers, Hilda Lau, Ila Marks, Eric Miller, Matthew J. Neave, Tobias A. Ross, Vittoria Stevens, Duncan Sutherland, Marcel Klaassen

## Abstract

The current panzootic of high pathogenicity avian influenza (HPAI) H5N1 has been catastrophic for wildlife, and following a significant sweep, clade 2.3.4.4b is found in every region aside from Oceania. Herein, we report the results of our third year of targeted surveillance of incoming migratory seabirds and shorebirds into Australia. We did not find evidence of HPAI H5N1 in any of the birds tested, and there were no reports of HPAI H5N1 in wildlife tested through other surveillance schemes in 2024. Unlike previous years, we detected a diversity of low pathogenicity avian influenza (LPAI) viruses in shorebirds. Through phylogenetic analysis we revealed that the H3N7 and H4N7 viruses recovered from Red-necked Stints were complex mosaic viruses, comprising segments of Eurasian, Australian shorebird, and Australian waterfowl segments. A H1N7 virus detected comprised a wholly Eurasian introduction, confirming this route for avian influenza viruses into Australian ecosystems. These results provide further evidence for the key role of long-distance migratory shorebirds in introducing novel LPAI viruses into Oceania. While our focus on northern migration routes remained appropriate for HPAI H5N1 surveillance in 2024, the continued spread of HPAI H5N1 to sub-Antarctic Islands demands consideration of a potential southern incursion route for Oceania in future.

## Introduction

Globally, the high pathogenicity avian influenza (HPAI) H5N1 panzootic is having an ongoing and significant impact on livestock and wildlife [*e.g.* 1-5]. While HPAI H5N1 has been circulating since 1996, with the start of the panzootic in 2021, clade 2.3.4.4b has caused a near global sweep, driving up the number of notifications associated with an enormous and rapid geographic and host range expansion [1-5]. Hundreds of millions of poultry have been culled [6] more than 400 species [1] of wild birds have been affected and millions have died [1,7], with some species facing population or species-level consequences [*e.g.* 2,8-10]. HPAI H5N1 has been detected in more than 50 mammalian species [4,5], with mammal-to-mammal transmission occurring in marine mammals in South America [11] and in dairy cattle in the USA [12,13]. More than 70 human cases have been confirmed since 2024 [14,15]. Overall, the arrival of HPAI H5N1 to new continents and into new animal species has been catastrophic.

HPAI H5N1 2.3.4.4b has now been detected in all regions of the world except Oceania [16]. The threat of HPAI virus arrival to the region is most likely to come from long-distance migratory birds, with recent evidence demonstrating that migratory shorebirds can import Asian-lineage low pathogenicity avian influenza (LPAI) H5 into Australia [17]. Millions of migratory shorebirds and seabirds arrive in Oceania each spring [18], creating an annual window of heightened incursion risk. There are no long-distance migratory waterfowl transiting between Australia and Asia, such that they are unlikely to contribute to virus incursion into Oceania (but may contribute to spread once the virus has arrived). This lack of migratory waterfowl is likely a key reason why HPAI has not arrived in Australia, despite being present in Asia since 1996 [16]. While seabirds are not traditionally implicated in the long-distance spread of HPAI, virus expansion in South America and Antarctica has been driven by seabirds [3,19]. While the Antarctic/sub-Antarctic route does provide a third avenue for introduction into Oceania, in 2024, outbreaks were still occurring many thousands of kilometres from the Australian mainland during 2024 (https://scar.org/library-data/avian-flu). All the while, HPAI outbreaks have been ongoing in south-east Asia [6], which are important stop-over sites for incoming migratory shorebirds into Australia.

Herein we report the results of our third year of targeted surveillance of incoming migratory seabirds and shorebirds into Australia [16,20]. Once again, we did not find evidence that these birds were infected with or had been exposed to HPAI H5N1; however, unlike previous years, we detected a diversity of LPAI viruses in shorebirds, all of which had at least one segment of Eurasian origin. This provides further evidence for the key role of shorebirds in introducing novel avian influenza genetic diversity into Australia, and that shorebird migration continues to be a potential pathway for incursion of HPAI H5N1.

## Methods

### Ethics statement

Capture, banding and sampling were conducted under ABBBS authorities 2824 to Tegan Douglas, 2915 to Marcel Klaassen and 8001 to the Victorian Wader Study Group, and approval of animal ethics committees of Deakin University (B28-2023), Department for Environment and Water South Australia Wildlife Ethics Committee (45/2022), Natural Resources and Environment Tasmania (AEC Project No. 6/2022-23 NRE), Department of Primary Industries and Regional Development Western Australia (WAEC 23-08-52) and Charles Sturt University (A22382).

### Sample collection and screening

We captured and sampled Short-tailed Shearwaters (*Ardenna tenuirostris*) at a breeding colony on Phillip Island, Victoria (38° 31′ 40″ S, 145° 20′ 13″ E) and Providence Petrel (*Pterodroma solandri*), Wedge-tailed Shearwater (*Ardenna pacifica*) and Flesh-footed Shearwaters (*hereafter* Sable Shearwater [21]) (*Ardenna carneipes*) at a breeding colony on Lord Howe Island, New South Wales (31° 33′ 15″ S, 159° 5′ 6″ E), upon their arrival from the northern Pacific. We also sampled 11 Asian-breeding migratory shorebird species at major nonbreeding sites in Western Australia (Exmouth Gulf; 22° 10′ 0″ S, 114° 18′ 0″ E) Victoria (Yallock Creek in Western Port Bay; 38° 22′ 0″ S, 145° 20′ 0″ E, Port Philip Bay, notably Western Treatment Plant; 38° 0′ 0″ S, 144° 34′ 0″ E, Port Fairy; 38° 22′ 0″ S, 142° 14′ 0″ E), South Australia (Limestone coast 37°56’57.4"S 140°28’04.1"E), and Tasmania (King Island; 39° 52′ 21″ S, 143° 59′ 8″ E) (Table 1, Figure 1). Migratory shorebirds were captured using cannon-netting and walk-in trapping, and seabirds were captured by picking them up from the ground in the colony.

**Figure 1:**
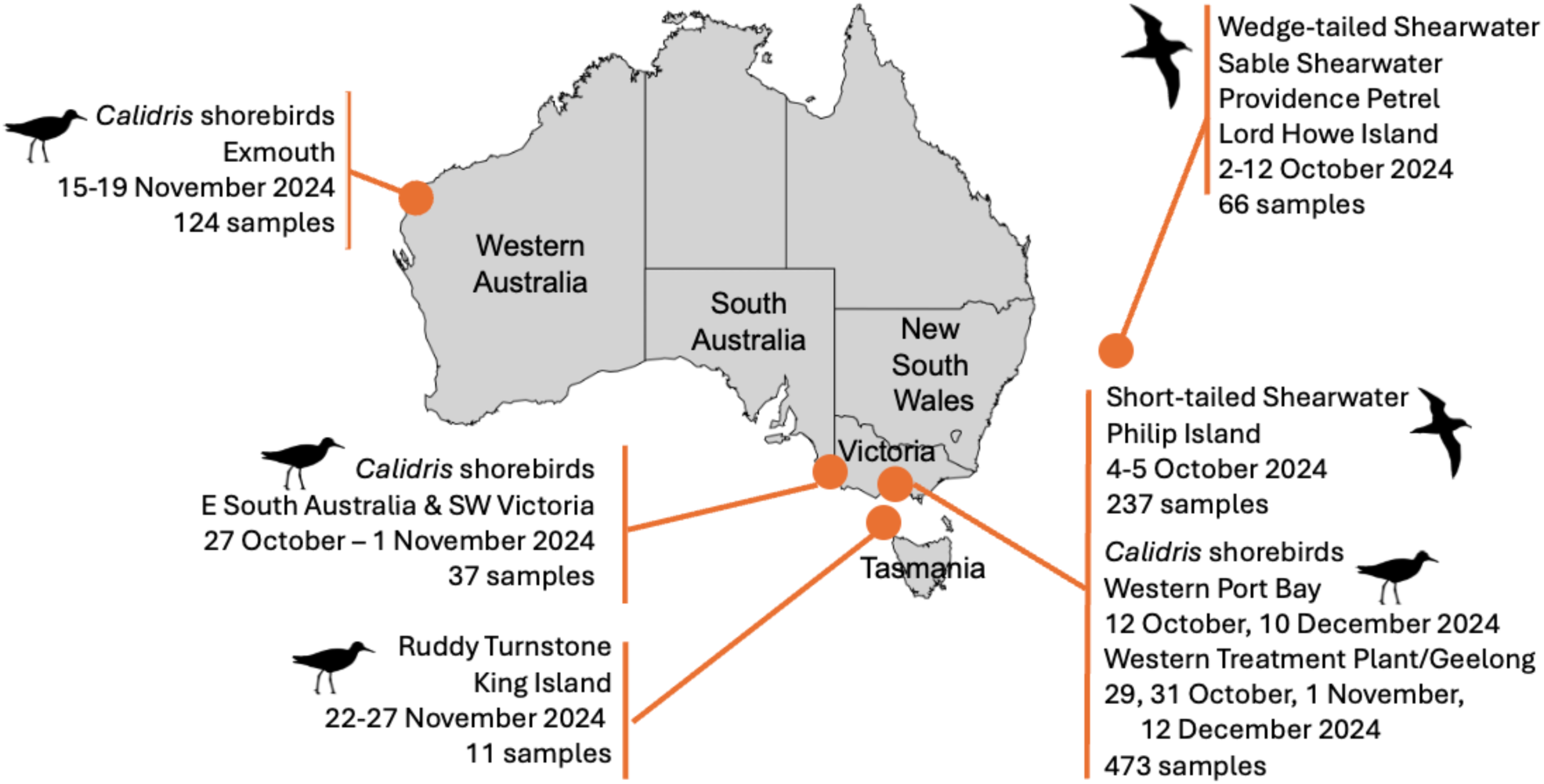
Overview of migratory bird sampling between October-December 2024 including locations, main target species, and number of samples collected.

**Table 1:** Migratory species targeted for active HPAI surveillance following arrival to Australia after migration from Asia and North America.

| Location | Dates | Species | Virus detection (qPCR) | Antibodies (NP-ELISA) |
| --- | --- | --- | --- | --- |
| Lord Howe Island, New South Wales | 2-12 October 2024 | Sable Shearwater<br>*Ardenna carneipes* | 0/37 | 2/35 |
|  |  | Providence Petrel<br>*Pterodroma solandri* | 0/6 | 0/6 |
|  |  | Wedge-tailed Shearwater<br>*Ardenna pacifica* | 0/23 | 0/22 |
| Philip Island, Victoria | 4-5 October 2024 | Short-tailed Shearwater<br>*Ardenna tenuirostris* | 0/237 | 28/235<br>(3 HI LPAI H5 positive) |
| Yallock Creek, Victoria | 12 October 2024 | Curlew Sandpiper<br>*Calidris ferruginea* | 0/3 | 0/3 |
|  |  | Little Stint<br>*Calidris minuta* | 0/1 | 0/1 |
|  |  | Red-necked Stint<br>*Calidris ruficollis* | 3/83 | 17/82 |
|  |  | Sharp-tailed Sandpiper<br>*Calidris acuminata* | 0/20 | 3/19 |
| Eastern South Australia & Port Fairy, Victoria | 27 October - 1 November 2024 | Ruddy Turnstone<br>*Arenaria interpres* | 0/14 | 3/14 |
|  |  | Red-necked Stint<br>*Calidris ruficollis* | 4/20 | 13/19 |
|  |  | Sanderling<br>*Calidris alba* | 0/3 | 0/3 |
| Port Philip Bay (notably Western Treatment Plant), Victoria | 29 October - 1 November | Red-necked Stint<br>*Calidris ruficollis* | 4/13 | 3/11 |
|  |  | Sharp-tailed Sandpiper<br>*Calidris acuminata* | 1/43 | 11/43 |
| Exmouth Gulf, Western Australia | 5-19 November 2024 | Bar-tailed Godwit<br>*Limosa lapponica* | 0/8 | 0/8 |
|  |  | Greater Sandplover<br>*Charadrius leschenaultii* | 0/32 | 0/32 |
|  |  | Common Greenshank<br>*Tringa nebularia* | 0/1 | 0/1 |
|  |  | Grey-tailed Tattler<br>*Tringa brevipes* | 0/10 | 0/10 |
|  |  | Lesser Sand Plover<br>*Charadrius mongolus* | 0/1 | 0/1 |
|  |  | Red-necked Stint<br>*Calidris ruficollis* | 0/37 | 10/36 |
|  |  | Ruddy Turnstone<br>*Arenaria interpres* | 0/32 | 3/32 |
|  |  | Sanderling<br>*Calidris alba* | 0/3 | 0/3 |
| King Island, Tasmania | 22-27 November 2024 | Red-necked Stint<br>*Calidris ruficollis* | 0/2 | 0/2 |
|  |  | Ruddy Turnstone<br>*Arenaria interpres* | 0/9 | 6/8 |
| Yallock Creek, Victoria | 10 December 2024 | Curlew Sandpiper<br>*Calidris ferruginea* | 0/13 | 0/13 |
|  |  | Red-necked Stint<br>*Calidris ruficollis* | 5/138<br>(1 H5 qPCR positive) | 36/138 |
|  |  | Sharp-tailed Sandpiper<br>*Calidris acuminata* | 0/12 | 1/12 |
| Western Treatment<br>Plant, Port Philip<br>Bay, Victoria | 12 December<br>2024 | Curlew Sandpiper<br>*Calidris ferruginea* | 0/25 | 0/24 |
|  |  | Red-necked Stint<br>*Calidris ruficollis* | 13/100<br>(1 H5 qPCR<br>indeterminate) | 41/91 |
|  |  | Sharp-tailed Sandpiper<br>*Calidris acuminata* | 0/22 | 3/21 |

From each individual, we took a combined oropharyngeal and cloacal swab with a sterile tipped applicator placed in virus transport media. Samples were kept at 4°C for up to 10 days, before being stored at -70°C. We also collected up to 200µl of blood from shorebirds, and 200µl-400µl from seabirds. Blood was collected from the brachial vein using the Microvette capillary system for serum collection (Sarstedt) and centrifuged ∼10 hours after collection to separate serum from the red blood cells.

### Virus and antibody screening

Viral screening and sequencing was undertaken as per Wille *et al*. (2022) [22]. Briefly, RNA was extracted from the combined oropharyngeal/cloacal sample using the NucleoMag Vet Kit (Scientifix) on the KingFisher Flex platform (Thermo Fisher Scientific). RNA was screened by reverse transcriptase real time PCR (RT-qPCR) for the influenza A virus matrix gene [23] using the SensiFAST Probe Lo-Rox qPCR Kit (Bioline), and a cycle threshold (Ct) cut-off of 40.

Serology was undertaken in alignment with Wille *et al*. (2019) [16]. Briefly, all samples were tested for antibodies against any influenza A virus using the Multi Screen Avian Influenza Virus Antibody Test Kit (IDEXX). An S/N threshold of <0.6 was used. Positive serum for which there was enough volume remaining were then assayed using haemagglutinin inhibition (HI) against a 2.3.4.4b candidate vaccine virus A/Astrakhan/3212/2020(H5N8) as well as an Australian lineage LPAI H5N3 Australian lineage virus A/duck/Victoria/0305-2/2012(H5N3). The 2.3.4.4b candidate vaccine virus was a 6:2 recombinant virus on an A/Puerto Rico/8/1934(H1N1)(PR8) backbone with the multibasic cleavage site removed.

### Virus characterisation

Positive RNA was referred to the Australian Centre for Disease Preparedness (ACDP) for confirmation, H5/H7 RT-qPCR screening using validated assays, and when possible, full genome sequencing. For samples collected at Yallock Creek (Western Port Bay) in October 2024, samples were initially referred to Agriculture Victoria for confirmation and H5/H7 RT-qPCR testing, prior to being sent to ACDP for sequencing. At present, ACDP utilises two different H5 RT-qPCR assays: a broadly detecting H5 RT-qPCR, and an H5 RT-qPCR designed to detect only the LPAI H5 lineage which was circulating in Australia prior to 2023 [see 24,25].

Avian influenza positive swab samples were inoculated into 11-day old embryonated chicken eggs via the allantoic route. Following incubation for 3 days, allantoic fluid was harvested and tested for agglutinating activity using an haemagglutination assay (HA). Isolate RNA was extracted using the QIAamp Viral RNA Mini Ki (Qiagen).

RNA from original samples was sequenced as per Wille *et al.* (2022) [24]. Briefly, up to 24 samples pooled per sequencing run by use of dual index library preparation and the Nextera XT DNA Library Preparation kit and 300-cycle MiSeq Reagent v2 kit (Illumina). Sequence reads were trimmed for quality and mapped to respective reference sequence for each influenza A virus gene segment using Geneious Prime software (www.geneious.com) (Biomatters, Auckland, NZ).

RNA extracted from viruses isolated in embryonated chicken eggs was sequenced using Oxford Nanopore Technologies (ONT). Specifically, amplicons for influenza A viruses were generated using previously published primers [26] with the SuperScript IV One- Step RT-PCR system (ThermoFisher Scientific). Fifty nanograms of normalized amplicons were used for library preparation using the ONT Rapid Barcoding Kit (Oxford Nanopore Technologies). Libraries were run on ONT R10 Standard Flow Cell (Oxford Nanopore Technologies) using a MinIon Mk1b for 8 hours. Nanopore reads were base-called using Dorado 7.4.14 within the MinKNOW package. Genomes were assembled using the IRMA pipeline [27].

### Phylogenetic analysis

All H1, H3, H4, N7 sequences (>1650bp for HA, and >1400bp for NA) from birds were downloaded from BV-BRC (07/01/2025). For internal genes, “global” trees were based on tree backbones from [17] and were supplemented with the top 20 blast hits from each sequence (07/01/2024). Sequences were aligned using MAFFT [28] integrated within Geneious Prime. Maximum likelihood trees incorporating the best-fit model of nucleotide substitution were estimated using IQ-Tree [29] with 1000 ultrafast bootstraps.

For segments and clades wherein the most closely related viruses were those from Australia, we downloaded the top 10 blast hits (downloaded via Geneious 28/01/2025). Sequences were aligned and maximum likelihood trees were constructed as above.

For segments and clades wherein the most closely related viruses were of Eurasian origin, we performed time scaled phylogenies. Trees were constructed using the top blast hits and included 10 sequences from 2010-2019 from the top 100 blast hits (30/01/2025) to ensure clock-like-behaviour. In cases where a clock-like structure was not achieved following this protocol, additional sequences from 2000-2010 were added. We evaluated the extent of molecular clock-like structure in the data by performing linear regressions of root-to-tip distances against year of sampling using maximum likelihood trees using TempEst [30]. Time scaled phylogenetic trees were estimated using BEAST v1.10.4 [31], under the uncorrelated lognormal relaxed clock [32] and bayesian skyline coalescent tree prior [33]. We used the SRD06 codon structured nucleotide substitution model [34], with the exception of M and NS segments for which we used the HKY+G model due to overlapping reading frames. One hundred million generations were performed, and convergence was assessed using Tracer v1.8 (http://tree.bio.ed.ac.uk/software/tracer/). Maximum credibility lineage trees were generated using TreeAnnotator following the removal of 10% burnin, and trees were visualised using Fig Tree v1.4 (http://tree.bio.ed.ac.uk/software/figtree/).

## Results

### Sample collection

To reveal whether a viral incursion may have occurred in Australia in 2024 with the arrival of wild migratory seabirds and shorebirds, we sampled 946 migratory birds of the order Charadriiformes and Procellariiformes, between October and December 2024. However, as we were not always successful in blood and swab sampling from each individual, a total of 925 serum samples and 948 swab samples were collected (Figure 1, Table 1).

### No confirmed HPAI H5 detected in migratory shorebirds in 2024

Overall, 30 samples were RT-qPCR positive for influenza A virus, of which 29/30 were from Red-necked Stints (*Calidris ruficollis*) collected in Victoria. One RT-qPCR positive was from a related species, Sharp-tailed Sandpiper (*Calidris acuminata*), also collected in Victoria. H5/H7 RT-qPCR testing revealed that all samples were negative for H7. One sample (sample ID: 20052) was positive for H5, and one sample (sample ID: 19814) was indeterminate (Ct >35) for H5. We were unable to resolve the H5 lineage using RT-qPCR assays due to the high Ct values. However, the most parsimonious conclusion is that we detected a LPAI H5. This conclusion is based on the observation that of the bird from which the sample was taken, as well as all birds caught and observed on the site at catch and since, were apparently healthy with no obvious signs of disease. Furthermore, in cases where wild bird mortality events occurred in Australia during this period, all tested birds were negative for HPAI [35].

### Seroprevalence in most migratory birds stable across multiple years

180 serum samples tested positive for anti-NP antibodies using a commercial ELISA (given an S/N cut-off of 0.6, as used previously [*e.g.* 16] (Table 1). In alignment with previous reports 16,20,22], antibodies were detected in Red-necked Stints, Ruddy Turnstones (*Arenaria interpres*), Sharp-tailed Sandpipers, Sable Shearwater and Short-tailed Shearwaters. Moreover, for all species seroprevalence also fell within previously reported ranges (*i.e.* not statistically significant based on 95% confidence intervals overlapping; Table 1, Figure S1). The exception is Red-necked Stints, for which seroprevalence was higher in 2024 (120/379=32%) relative to previous years. This is likely related to the high LPAI prevalence (reported here) detected in this species compared to previous years wherein we didn’t detect any positive samples during the same time of year [16,20].

The 129 samples positive by anti-NP-ELISA that had sufficient sample volume remaining were assayed by haemagglutination inhibition (HI) using a lineage 2.3.4.4b virus and an Australian lineage LPAI H5N3 virus. No birds demonstrated serological evidence of exposure to HPAI H5 2.3.4.4b. However, three Short-tailed Shearwater had antibodies against the LPAI H5 Australian lineage virus.

### Novel incursions of LPAI detected in shorebirds in 2024

Whole-genome sequencing resolved the subtypes of 16 viruses, comprising H3N7 (n=8), H4N7 (n=6), H1N7 (n=1) and a mixed H3/H4N7 (n=1). All genomes were recovered from samples collected from Red-necked Stints sampled in Victoria, with the exception of an H1N7 virus from a Sharp-tailed Sandpiper also sampled in Victoria (Table 2). The sample genotypes of both H3N7 and H4N7 viral genomes were recovered from sampling events in October and December from two sampling sites (Yallock Creek and Western Treatment Plant), which are within 100km, suggestion transmission in the local shorebird population over a two month period (Table 2).

**Table 2:**
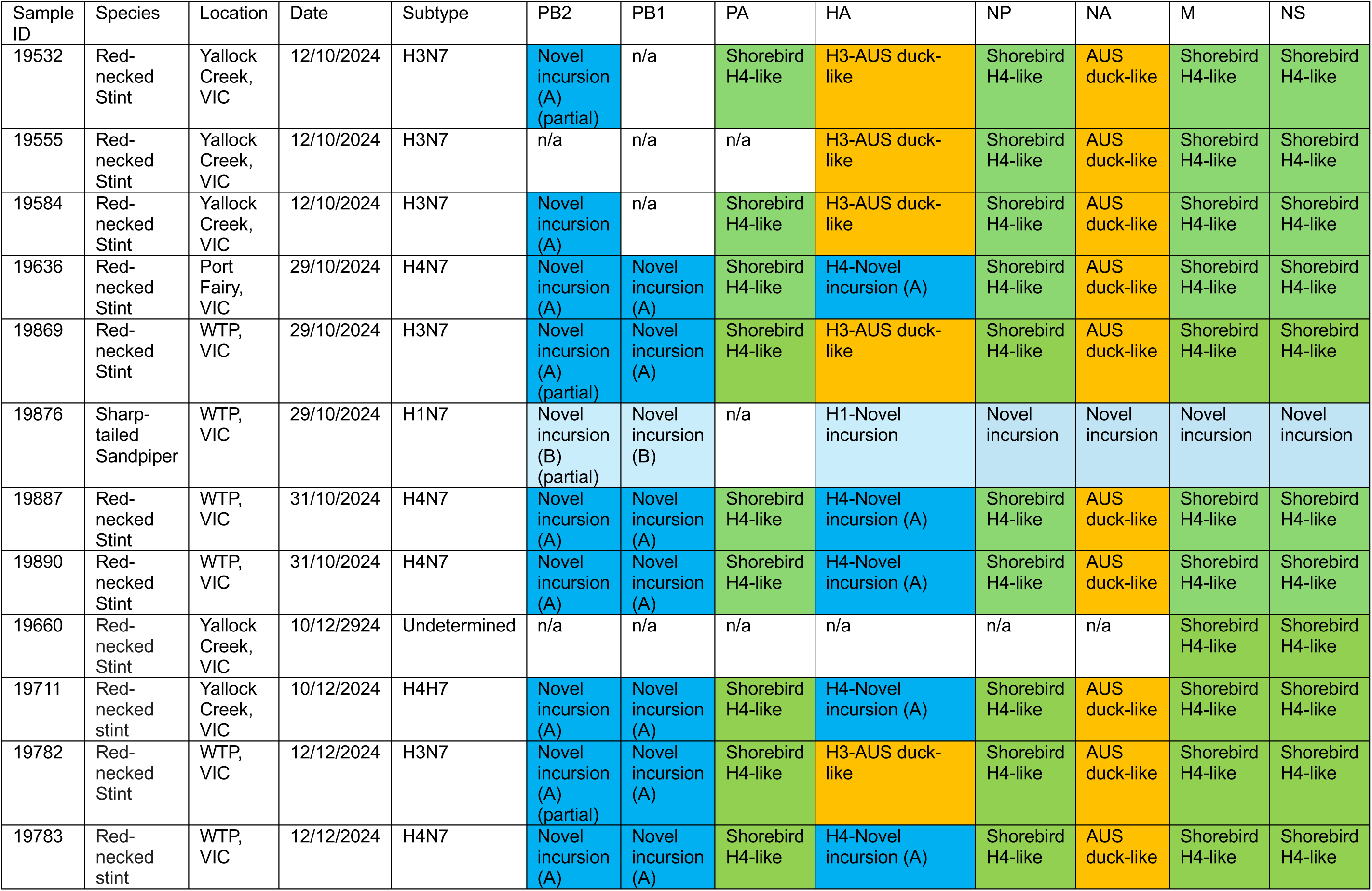
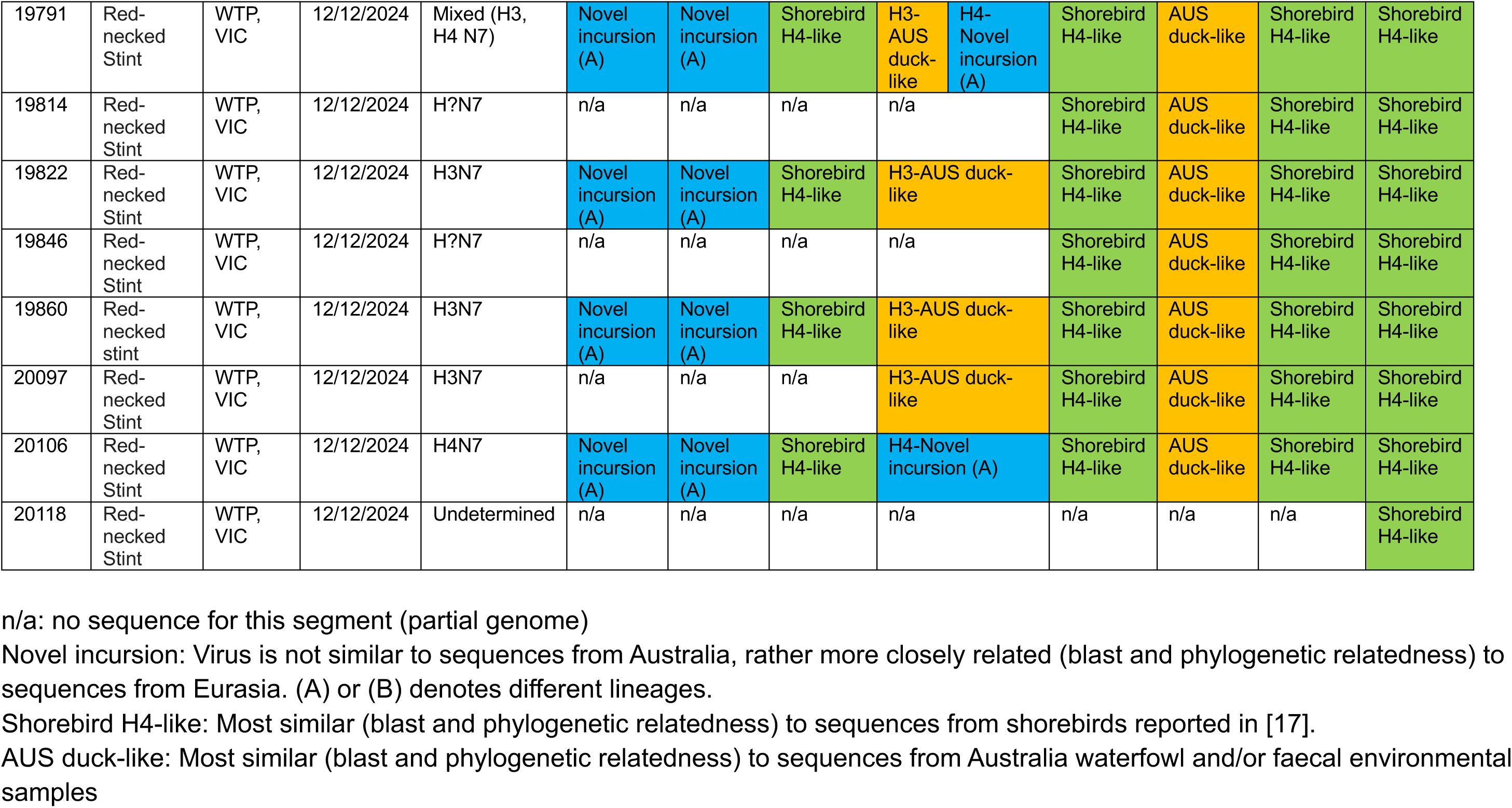
Genome constellations of LPAI viruses sequenced in this study.

Of these, we generated 11 complete genomes, and nine partial genomes (Table 2). A diversity of genome constellations (genotypes) were recovered. All genomes had at least one segment comprising a novel incursion into Australia (*i.e.* more similar to sequences from Eurasia than from Australia), of which all recovered PB2 and PB1 segments (regardless of subtype) comprised Eurasian sequences. The eight H3N7 genomes were comprised of two Eurasian segments (PB2, PB1), whereas the H4N7 viruses had three Eurasian segments (PB2 PB1, HA), and the H1N7 virus was entirely of Eurasian origin (7/7 recovered segments, we did not recover PA) (Table 2, Figure 2-3, Figure S2-S4).

**Figure 2.**
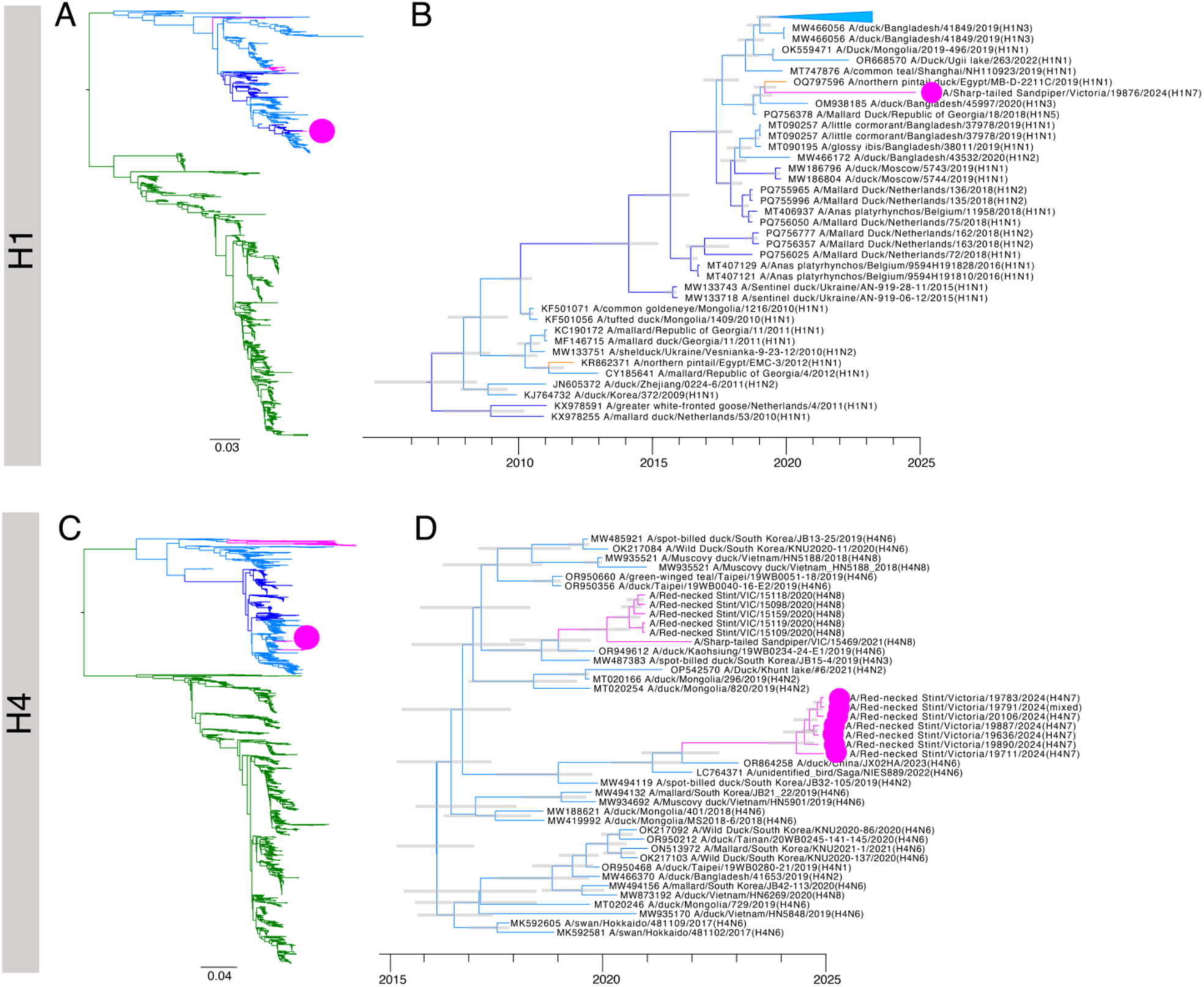
Phylogenetic trees of (a–b) H1 and (c-d) H4 sequences. (a and c) Maximum likelihood trees comprising all sequences collated for this study. Trees were rooted geographically (i.e. between the ‘Eurasian’ and ‘American’ lineages), and the scale bar corresponds to the number of substitutions per site. (b and d) Time-structured phylogenetic trees. The trees comprise select sequences from blast hits. Node bars correspond to the 95% HPD of node height. Sequences of interest are indicated by arrows in (a and d) or a filled circle. Branches are coloured by continent.

**Figure 3.**
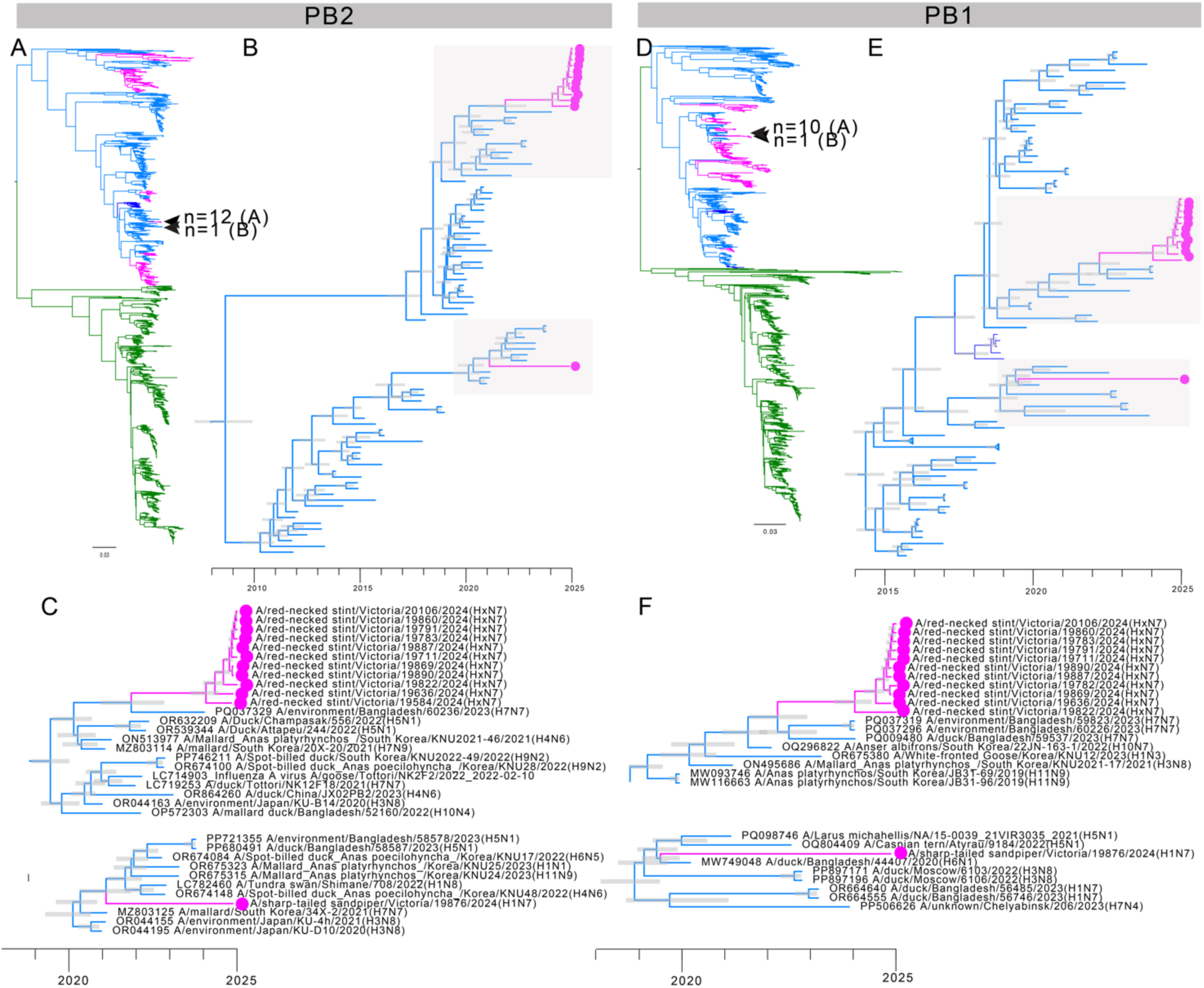
Phylogenetic trees of (a–c) PB2 and (d–f) PB1 sequences. (a and d) Maximum likelihood trees comprising all sequences collated for this study. Trees were rooted geographically (i.e. between the ‘Eurasian’ and ‘American’ lineages), and the scale bar corresponds to the number of substitutions per site. (b and e) Time-structured phylogenetic trees. The trees comprise select sequences from blast hits. Node bars correspond to the 95% HPD of node height. (c and f) Expansion of clade from b and e, which have been highlighted in a grey box. Sequences of interest are indicated by arrows in (a and d) or a filled circle in b, c, e, f. Branches are coloured by continent.

The H3 HA segment was most closely related to viruses recovered from wild waterfowl in Australia (98% similar to top blast hit OL371991 A/wild waterfowl/Queensland/P19-01624-17/2019(H3N8)), and fell into a lineage that has been circulating in Australian waterfowl for decades (Figure S2). In contrast, the H4 HA sequences were most closely related to sequences from Eurasia, including sequences from China and South Korea (98.5% similar to OR864258 A/duck/China/JX02HA/2023(H4N6)) (Figure 2). There was approximately three years between the mean most recent common ancestor (MRCA) of the sequences recovered in this study (Sept 2021, 95% highest posterior density (HPD) Nov 2020-Aug 2022), and date of divergence from the closest sequence in GenBank (April 2024, 95% HPD Sept 2023-Sept 2024), making inferences around viral incursion challenging (Table S1, Figure S5). All H3 and H4 viruses shared the same N7 sequence (mean % identity 99.2%), which fell into a clade of Australian sequences (98.5% similar to closest blast hit OL370323 A/wild duck/New South Wales/M19-11593-66/2019(H7N7)) which has been circulating in Australia since ∼2016 [24] (Table 2, Figure S2).

The H3 and H4 viruses comprised complex reassortants, sharing segments with origins from viruses in Eurasia, Australian shorebirds, and Australian waterfowl (Table 2). Time-scaled phylogenetic methods revealed that the date of divergence from the closest GenBank sequence and the MRCA of the PB1 and PB2 (shared by H3 and H4 viruses, Figure 3) and H4 HA (Figure 2) overlapped (Table S1, Figure S5). This suggests that the H4, PB1, and PB2 segments (of H3 and H4 viruses) likely arrived simultaneously. Following arrival in Australia, a series of reassortment events likely occurred. First, the PA, NP, M, NS segments of both the H3 and H4 viruses were most closely related to H4N8 viruses recovered from Victorian shorebirds in 2020 and 2021 [17] (Table 2, Figure S3). This lineage has most likely been circulating in shorebirds since ∼2012, with detections predominantly in Victorian Red-necked Stints (detected in 2012, 2017, 2020), but also in Red-necked Stints sampled in Japan (2012) [17]. The N7 of both H3 and H4 was most closely related to Australian waterfowl, as was the H3 HA, (Table 2, Figure S2) suggesting a further reassortment event having occurred with viruses from Australian waterfowl. The intricate details of these local reassortment events cannot easily be resolved given the patchy sequence data, and the long-time intervals as revealed by time structured analysis.

Blast and phylogenetic analysis revealed that the H1 HA sequence was most closely related to sequences from Egypt (97.7% similar to OQ797596 A/northern pintail duck/Egypt/MB-D-2211C/2019(H1N1)) Republic of Georgia (98.1% similar to PQ756378 A/Mallard Duck/Republic of Georgia/18/2018(H1N5)) and Europe (e.g. 97.7% similar to PV738505 A/wild goose/Germany-HE/2025AI02191/2025(H1N2)) (Figure 2). Time-structured analysis demonstrated that the date of divergence from the closest GenBank sequence was February 2019 (95% HPD Sept 2018-July 2019) (Table S1). The 5-year interval between the detection in Australia and date of divergence from the closest GenBank sequence makes it extremely challenging to infer where this clade was circulating prior to detection in Australia, and what the exact time window of introduction into Australia was. While this H1N7 genome shared the same NA subtype as the H3/H4 viruses, the N7 sequence was dissimilar (only 90.9% similar to the N7 sequences from the H3/H4 genomes) (Figure S4). Rather, the N7 sequence is most similar to a virus from Bangladesh (e.g. 98.87% similar t0PQ037279 A/environment/Bangladesh/ 59969/2023(H7N7)), with the mean MRCA of the clade containing the N7 sequence generated here and the most closely related sequence being Oct 2022 (95% HPD Dec 2021-June 2023) (Figure S4). Unlike large number of shared segments between the H3 and H4 viruses, all segments for the H1N7 virus were unique and were unrelated to any Australian sequences (with an average of only 92% similarity to the H3/H4 segments recovered). The most closely related sequences in GenBank were of viruses from wild birds in Asia (Korea, Bangladesh), and for the NS segment, Europe (98.57% similar to MK192309 A/mallard/Netherlands/25/2013(mixed)) (Figure 3, Figure S4). Like the HA segment, most of the segments of the H1N7 genome had a large time gap between the mean date of divergence from closest GenBank sequence (2014-2022) and date of detection (Table S1). Overall, this H1N7 virus likely comprises a wholly Eurasian introduction, or, while unlikely, a virus circulating in an unsampled reservoir in Australia.

## Discussion

Herein, we report the outcomes of the third year [16,20] of targeted surveillance undertaken in Australia to rule out a putative incursion of HPAI H5N1. Despite massive, global expansion of HPAI H5N1 since 2021 3,36,37], Oceania remains free of HPAI H5N1 [20]. In 2024, as in 2022 and 2023, there was no evidence of incursion of HPAI H5N1, which might be attributed to the limited movement of waterfowl between Australia and the rest of the world, despite the arrival to Australia of millions of individual migratory birds from other species. Waterfowl have been implicated as an important disperser of HPAI H5N1 given there is evidence that some species (e.g. Mallards *Anas platyrhynchos*) have no or limited clinical disease [38,39] and have been demonstrated to migrate whilst infected with HPAI H5N1 virus [40,41]. Importantly, however, with the exception of Australian waterfowl venturing into the northern parts of the Australo-Papuan region and vice versa, there are no waterfowl species migrating between Australia and Asia, the latter being a hotspot for HPAI H5N1 [18].

While the role of shorebirds as long-distance movers of HPAI H5N1 is unclear, they have been impacted by outbreaks [42,43], and have been demonstrated to survive HPAI H5N1 infection [44-46]. Furthermore, we provide additional evidence of their role as long distance movers of LPAI viruses. While geographic mosaic viruses have been previously described in shorebirds sampled in Australia [*e.g*. 17,47], we have potentially detected the first wholly Eurasian virus (H1N7) in Australia in a long-distance migrant. Moreover, the mosaic genome constellations of the H3 and H4 viruses sequenced in this study demonstrate reassortment between viruses previously only described in shorebirds [as reported in [17], and those from Australian waterfowl populations, demonstrating that viruses are being shared between shorebird and waterfowl hosts [48].

Australia’s surveillance approach for HPAI H5N1 considers northern routes of introduction given HPAI H5N1 has been circulating in Asia for decades. Notably, recent data from countries where robust surveillance systems are in place, like Japan and South Korea, confirm the frequent presence of HPAI H5N1 2.3.4.4b in wild birds in countries along the East Asian-Australasian Flyway [1]. Asia is unique in the co-circulation of multiple lineages of Goose/Guandong lineage HPAI H5Nx, a result of endemic circulation and diversification in poultry. Indeed, India, Bangladesh, Cambodia, and Indonesia report predominantly 2.3.2.1a or 2.3.2.1e in poultry and associated human cases, in addition to 2.3.4.4b [36,49-52]. However, unlike clade 2.3.4.4b, which is spreading with wild birds, clade 2.3.2.1a and 2.3.2.1e remain more restricted to poultry. It is possible that the ongoing presence of these other lineages of HPAI H5Nx may potentially result in cross-protection against 2.3.4.4.b in poultry, serving as a potential explanation why outbreaks of lineage 2.3.4.4b HPAI are remarkably limited in East Asia.

While our focus on northern migration routes remained appropriate for 2022-2024, the considerable spread of HPAI H5N1 2.3.4.4b in the Antarctic [19] requires the consideration of a potential southern incursion route. In South America and potentially the sub-Antarctic, seabirds and marine mammals appeared to contribute significantly to spread, deviating from the dogma that ducks are the main contributors to virus spread [3,19,53,54]. Virus genomes demonstrate that these outbreaks were caused by the spread of HPAI H5N1 2.3.4.4b from South Georgia Island, a distance of >5000km along the Southern Ocean to the French sub-Antarctic Islands [19]. Australia’s Heard and McDonald Islands are only 500km from Kerguelen Island, and seabirds breeding on Australia’s sub-Antarctic Islands (*e.g.* Macquarie Island) such as Northern Giant Petrels (*Macronectes halli*) travel along the easterly winds through this area [55]. While species implicated in the spread of HPAI H5N1 2.3.4.4b within the Antarctic region are only rarely seen in Australia, there are other seabirds breeding in Australia, such as the Short-tailed Shearwater, which we abundantly sampled (n=237), that may venture into Antarctic waters to forage [56] creating opportunities for species interactions and virus transmission. As such, it is critical for future surveillance to consider both an incursion from the north and from the south, by shorebirds and seabirds, respectively. Based on the timing of migration, an incursion from the north is most likely to occur during spring with the arrival of long-distance migrants. Providing a time window where incursion risk from the south is highest is more challenging given great uncertainties on what the potential (indirect) routes of entry may be.

## Acknowledgements

We are grateful to all those who assisted with sample collection in the field, specifically Alessia Ostolani (Linneaus University), Josefina Gutierrez (Universidad Austral de Chile), Sally Salmon (Agriculture Victoria), and all the volunteers of the Victorian Wader Study Group, Friends of Shorebirds SE, Victorian Ornithological Research Group, Philip Island Nature Parks. We are grateful to P. Eden and S. Ban (Wildlife Health Australia), and A. Breed (Australian Department of Agriculture, Fisheries and Forestry) for ongoing support.

## Data Accessibility

Sequences generated in this study have been deposited in GenBank (Accession numbers XXX). Phylogeny files have been made available on GitHub (https://github.com/michellewille2/EnhancedSurveillance2024/)

## Supplemental Tables and Figures

**Figure S1.**
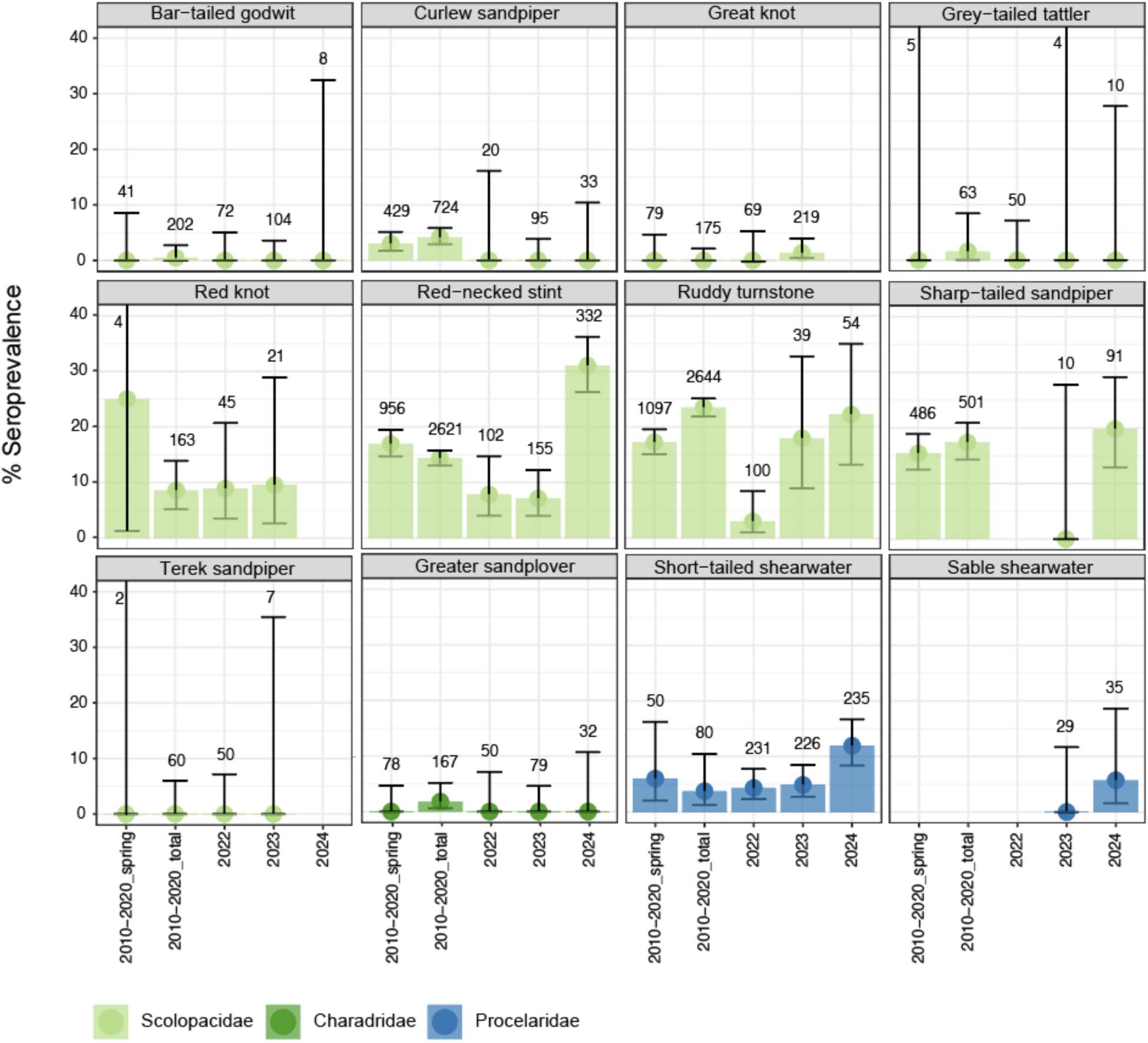
No change in seroprevalence in migratory species. For each species, we have shown prevalence data from: Wille et al. (2022), including data from Sept-Dec which is in alignment with the time window of this study, data from Wille et al. (2023), (d) data from Wille et al (2024), and finally, data presented in Table 1. Y axis comprises percentage seroprevalence, with 95% confidence intervals. In cases where the upper 95% interval extends beyond 40%, the upper has not been shown. Colours by avian family. Values comprise sample size. Species which were only sampled in one year have been excluded, and those with small sample sizes (<10) over the last three years have also been excluded. In cases where 95% confidence intervals overlap, the seroprevalence is not significantly different. For Red-necked Stint, 2024 is a clear outlier with higher seroprevalence compared with previous years (i.e. non-overlapping 95% confidence intervals).

**Figure S2.**
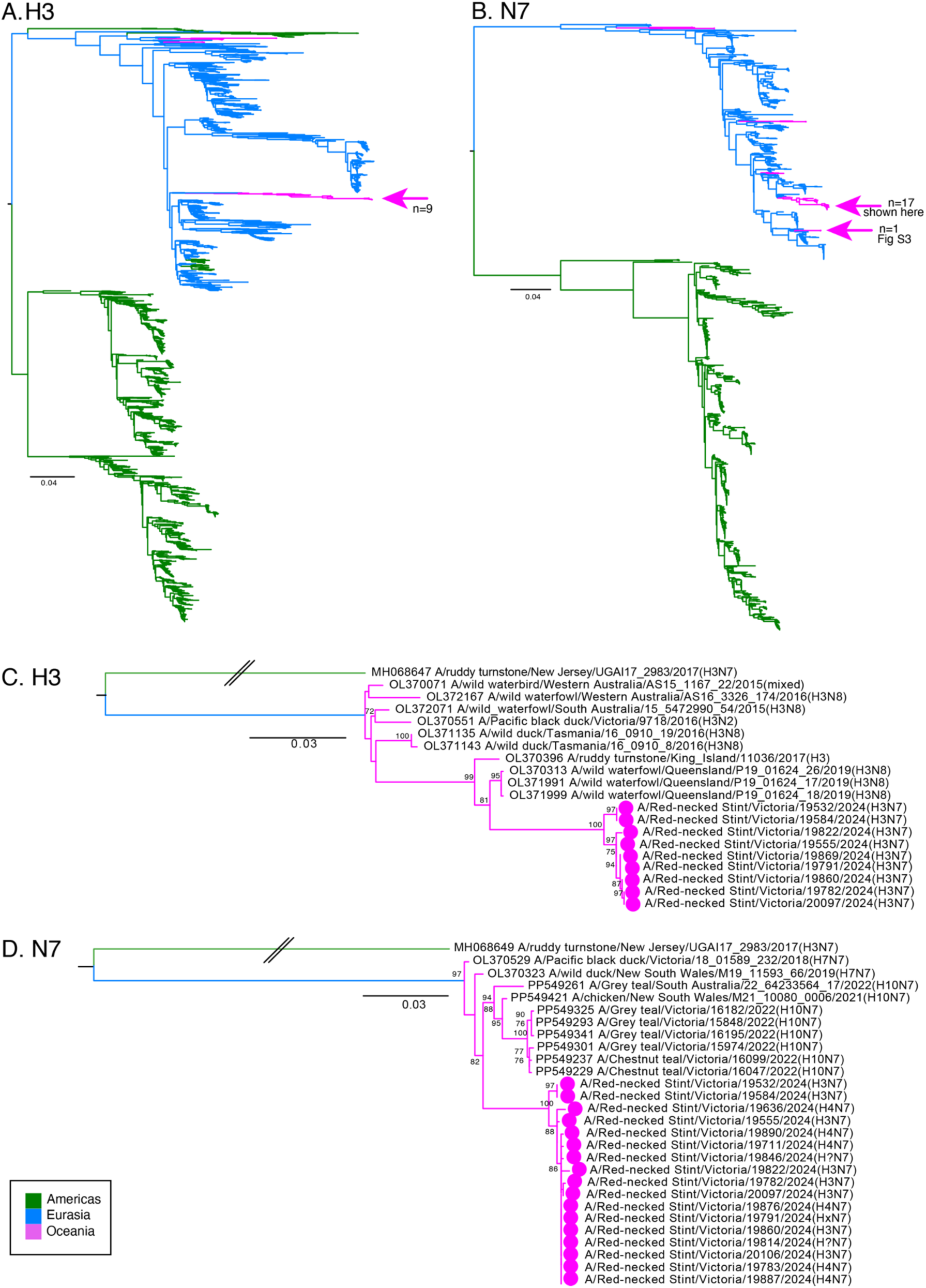
Phylogenetic trees for H3-HA and N7-NA sequences. (A, B) Maximum likelihood trees comprising all sequences from GenBank. Trees were rooted geographically (i.e. between the ‘Eurasian’ and ‘American’ lineages). Arrows correspond to sequences generated in this study (C,D) Maximum likelihood trees comprising the top 10 BLAST hits for H3 and N7 sequences which fall into established Australian clades. For the N7 of 19876, which is more similar to sequences from Eurasia, the corresponding phylogeny can be found in Figure S4. Trees rooted against A/ruddy turnstone/New Jersey/UGA17-2983/2017(H3N7) which falls into the “North American clade”. Scar bar corresponds to the number of substitutions per side. Node labels are ultrafast bootstrap values. Branches are coloured by continent

**Figure S3.**
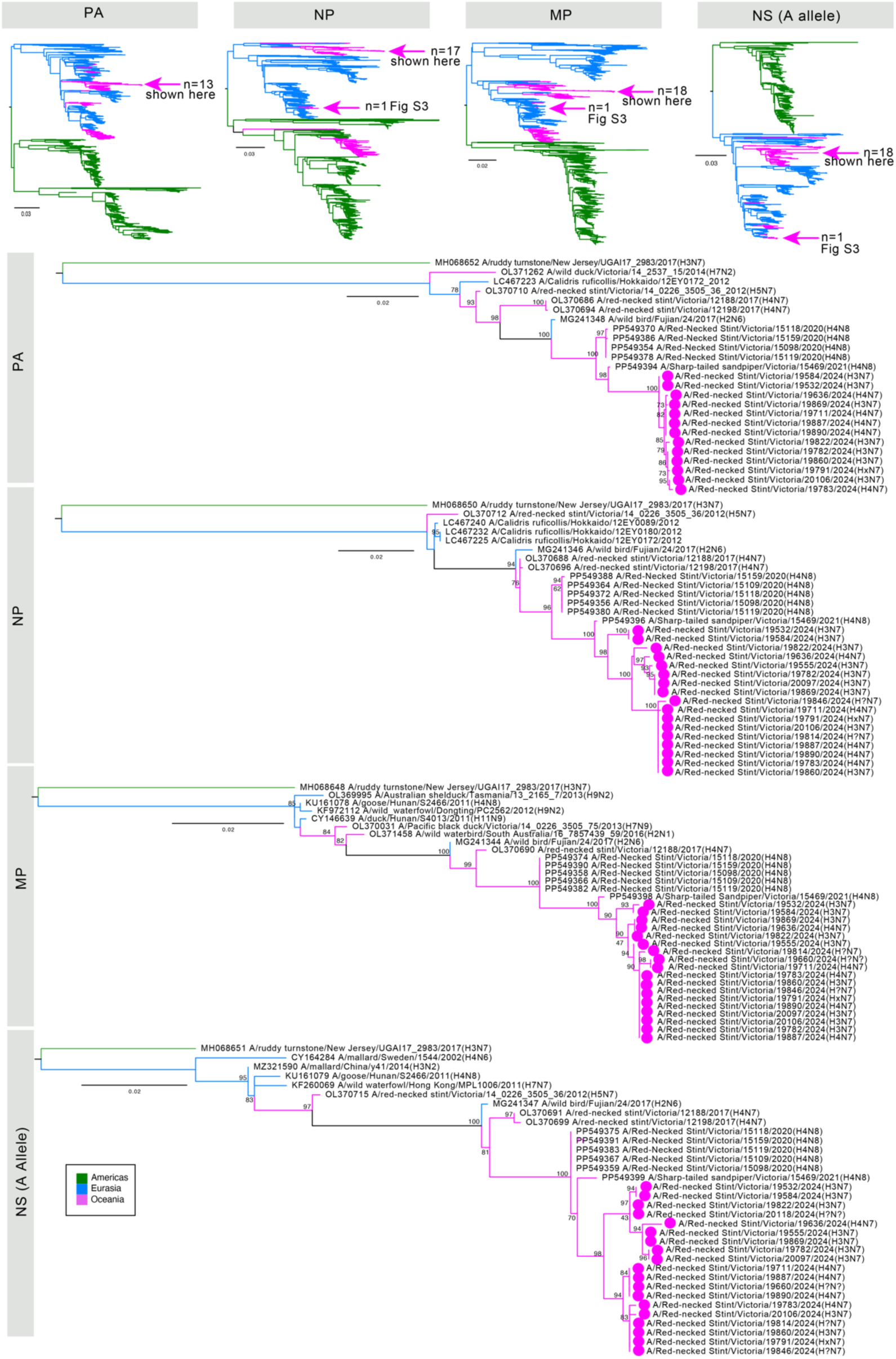
Phylogenetic trees for PA, NP, M, NS segments. (Upper) Maximum likelihood trees comprising representative global sequences (backbones from Wille et al. 2024). Trees were rooted geographically (i.e. between the ‘Eurasian’ and ‘American’ lineages). Arrows correspond to sequences generated in this study (Lower) Maximum likelihood trees comprising the top 10 BLAST hits sequences which fall into Australian clades, with the most closely related viruses those from shorebirds reported in Wille et al. 2024. For 19876, which is more similar to sequences from Eurasia, the corresponding phylogeny can be found in Figure S4. PB2 and PB1 trees can be found in Figure 3. Trees rooted against A/ruddy turnstone/New Jersey/UGA17-2983/2017(H3N7) which falls into the “North American clade”. Scar bar corresponds to the number of substitutions per side. Node labels are ultrafast bootstrap values. Branches are coloured by continent

**Figure S4.**
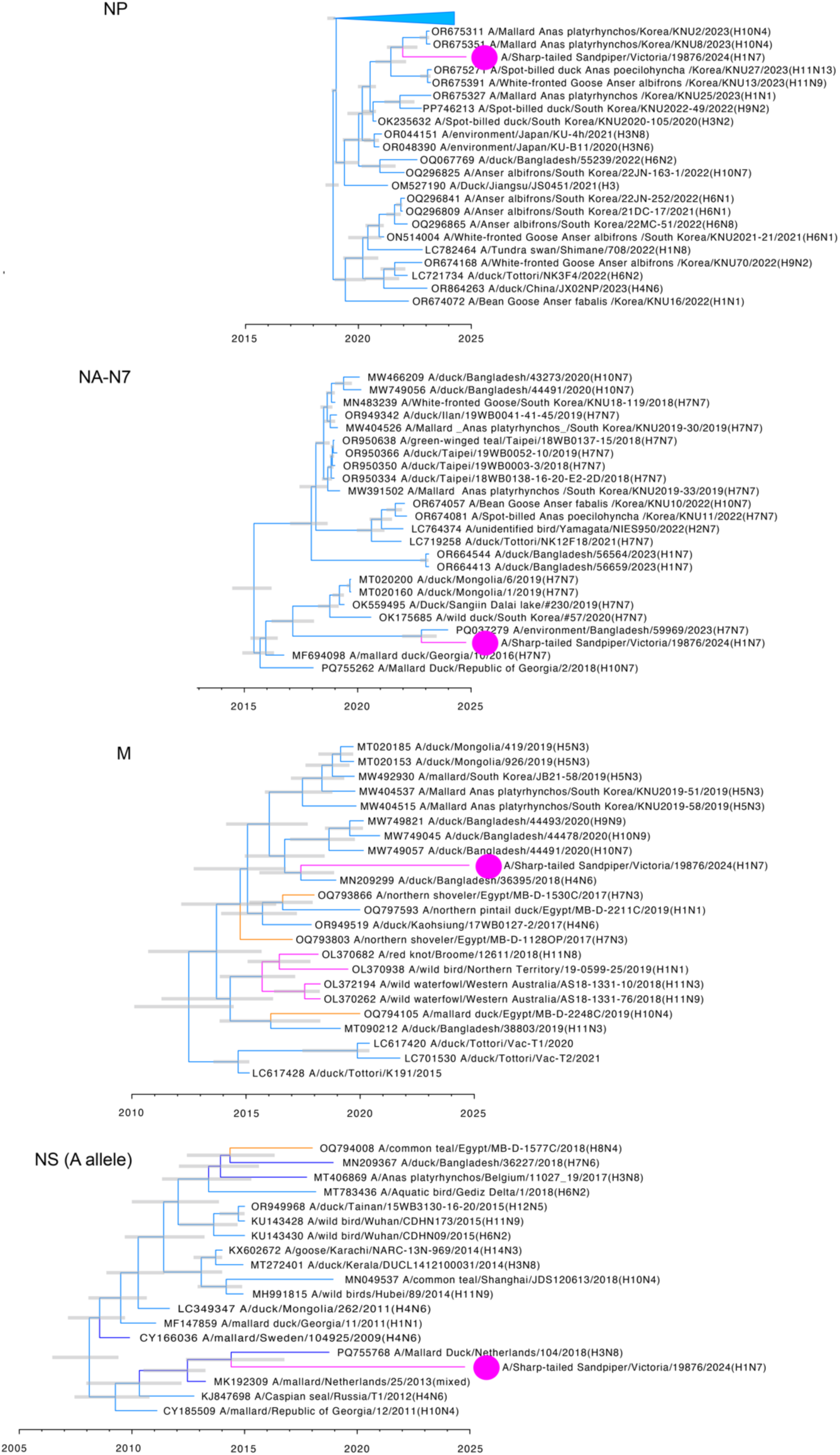
Phylogenetic trees of (A) NP, (B) N7, (C) M, (D) NS for A/Sharp-tailed Sandpiper/Victoria/19876/2024(H1N7). Node bars correspond to the 95% HPD of node height, scale bar is in years, and branches are coloured by continent. Maximum likelihood trees demonstrating placement in the “global” diversity of viruses for NP, M, and NS and can be found in Figure S3, and for N7 in Figure S2. Phylogenies for PB2 and PB1 can be found in Figure 3. As we did not recover the PA segment, there is no associated phylogenetic tree.

**Table S1.**
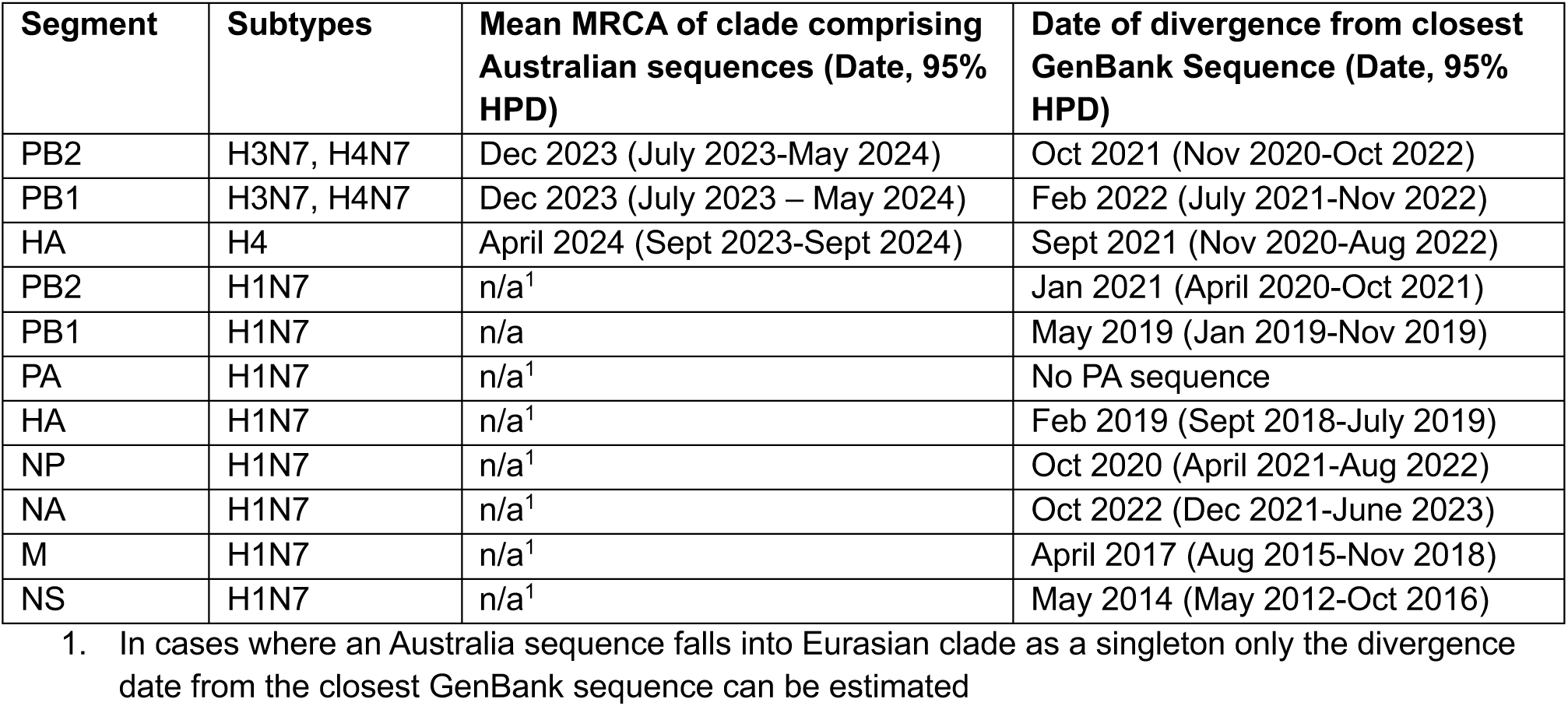
Estimated date of viral incursions into Australia for select segments and clades.

| Segment | Subtypes | Mean MRCA of clade comprising Australian sequences (Date, 95% HPD) | Date of divergence from closest GenBank Sequence (Date, 95% HPD) |
| --- | --- | --- | --- |
| PB2 | H3N7, H4N7 | Dec 2023 (July 2023-May 2024) | Oct 2021 (Nov 2020-Oct 2022) |
| PB1 | H3N7, H4N7 | Dec 2023 (July 2023 – May 2024) | Feb 2022 (July 2021-Nov 2022) |
| HA | H4 | April 2024 (Sept 2023-Sept 2024) | Sept 2021 (Nov 2020-Aug 2022) |
| PB2 | H1N7 | n/a<sup>1</sup> | Jan 2021 (April 2020-Oct 2021) |
| PB1 | H1N7 | n/a | May 2019 (Jan 2019-Nov 2019) |
| PA | H1N7 | n/a<sup>1</sup> | No PA sequence |
| HA | H1N7 | n/a<sup>1</sup> | Feb 2019 (Sept 2018-July 2019) |
| NP | H1N7 | n/a<sup>1</sup> | Oct 2020 (April 2021-Aug 2022) |
| NA | H1N7 | n/a<sup>1</sup> | Oct 2022 (Dec 2021-June 2023) |
| M | H1N7 | n/a<sup>1</sup> | April 2017 (Aug 2015-Nov 2018) |
| NS | H1N7 | n/a<sup>1</sup> | May 2014 (May 2012-Oct 2016) |

1. In cases where an Australia sequence falls into Eurasian clade as a singleton only the divergence date from the closest GenBank sequence can be estimated

**Figure S5.**
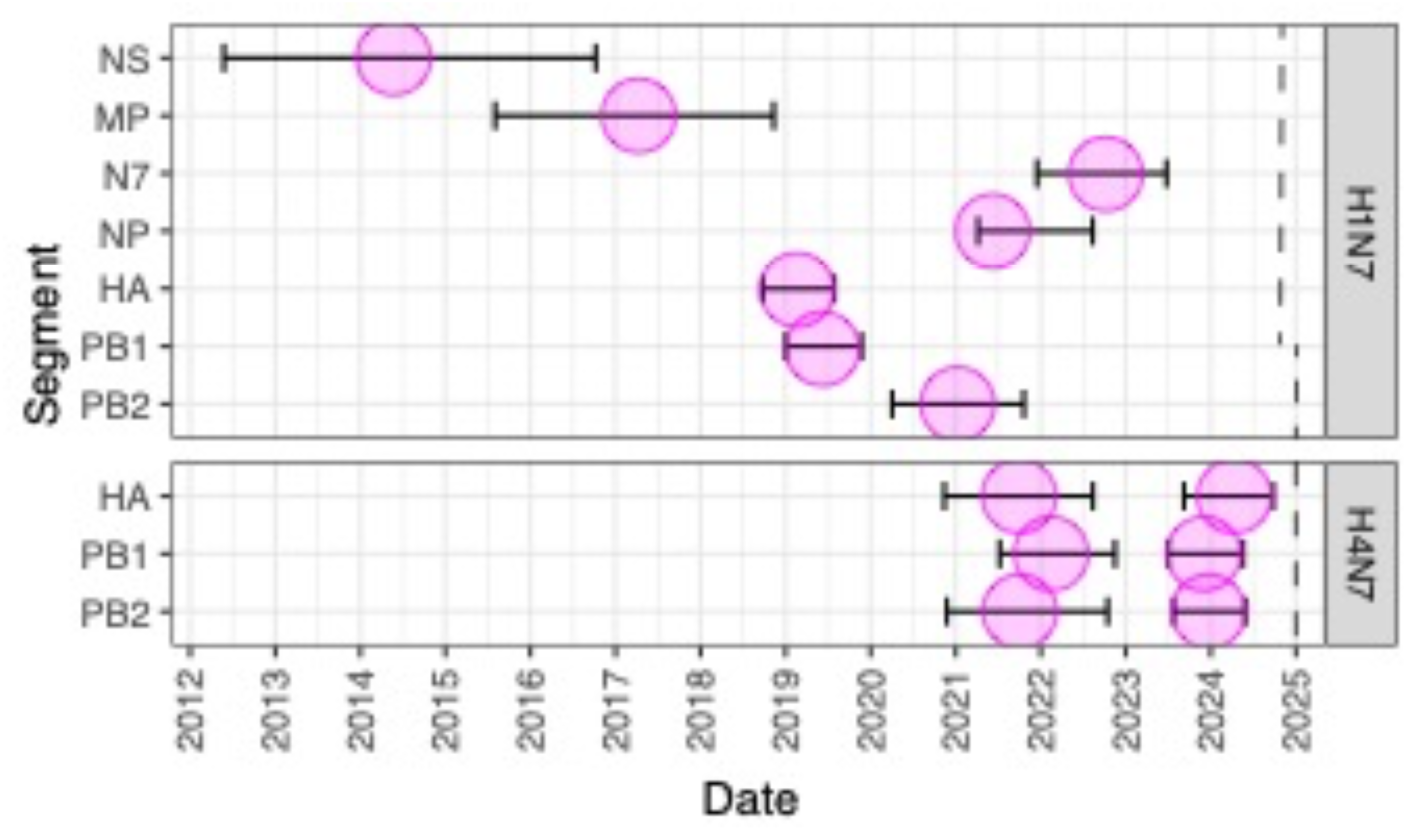
Estimated date of viral incursions into Australia for select segments and clades. For H1N7, we have only plotted the Date of divergence from closest GenBank Sequence. For H4N7 we have plotted both the Date of divergence from closest GenBank Sequence and the MRCA of the Australian clade for segments that comprise likely new viral incursions (as opposed to those in established Australian clades). Dotted line is the date of youngest sequence in the phylogenetic tree. Circle is the estimated mean MRCA, and the bars are the 95% HPD. Values in Table S1.

